# Towards Reduced Order Models via Robust Proper Orthogonal Decomposition to Capture Personalised Aortic Haemodynamics

**DOI:** 10.1101/2023.01.21.524933

**Authors:** Chotirawee Chatpattanasiri, Gaia Franzetti, Mirko Bonfanti, Vanessa Diaz-Zuccarini, Stavroula Balabani

## Abstract

Data driven, reduced order modelling has shown promise in tackling the challenges associated with computational and experimental hemodynamic models. In this work, we explore the use of Reduced Order Models (ROMs) to capture the main flow features in a patient-specific dissected aorta. We apply Proper Orthogonal Decomposition (POD) and Robust Principle Component Analysis (RPCA) on in vitro, hemodynamic data acquired by Particle Image Velocimetry and compare the decomposed flows to those derived from Computational Fluid Dynamics (CFD) data for the same geometry and flow conditions. The flow is reconstructed using different numbers of POD modes and the flow features obtained throughout the cardiac cycle are compared to the original Full Order Models (FOMs).

RPCA has been found to enhance the quality of PIV data and to capture most of the kinetic energy of the flow in just two modes similar to the numerical data that are free from measurement noise. The reconstruction errors differ along the cardiac cycle with diastolic flows requiring more modes for accurate reconstruction. In general, modes **Φ**1-10 are found sufficient to represent the flow field. The results demonstrate that the coherent structures that characterise this aortic dissection flow are described by the first few POD modes suggesting that it is possible to represent the macroscale behaviour of aortic flow in a low-dimensional space; thus significantly simplifying the problem, and allowing for more computationally efficient flow simulations that can pave the way for translation of such models to the clinic.

## 1. Introduction

The ability to visualize and investigate the flow features inside the aorta, either by *in vitro* experiments or Computational Fluid Dynamics (CFD), can provide invaluable information for clinical support, disease progression predictions and surgical treatment planning [1]. Application of such tools has been successfully demonstrated in several pathologies, such as aortic dissection [1, 2, 3], coronary artery disease [4], valve prosthesis [5], aortic aneurysm [6] and congenital heart disease [7]. Both in vitro and in silico hemodynamic approaches are subject to certain limitations that impact their clinical translation. In CFD for example, a compromise between model accuracy and complexity has often need to be made [8]. Over-simplifications of the geometrical domain and boundary conditions can lead to non-realistic results. On the other hand, increasing the model complexity further complicates the solution, increasing the computational time and often introducing or increasing parameter uncertainty. High computational cost represents a problem for the clinical translation of these numerical models, especially when considering the time-scales of *acute* pathological stages (i.e. days rather than weeks or months) and the limited time available in clinics to make full use of CFD as realistic tool for pre-interventional planning.

To address this problem, Reduced Order Models (ROMs) have been studied intensively in the last decade to enable faster calculations of fluid dynamic problems [9]. The main idea is to replace the large-scale problems by less complex ones, which can be solved with significantly less time and resources while maintaining an acceptable level of accuracy. Many methods have been developed to extract ROMs from high-fidelity data, such as Proper Orthogonal Decomposition (POD) [10, 11, 12], Proper Generalized Decomposition (PGD) [13], Dynamic Mode Decomposition [12, 14], Krylov subspace [15], Matrix interpolation [16], and the recently developed neural network based method, Autoencoder [17, 18, 19].

Among these methods, POD is arguably the most commonly used ^1^. POD provides a powerful tool to reduce the dimensionality of a system by projecting it to a set of Reduced Basis, called POD modes, which are orthogonal to each other. POD identifies the energetically dominant modes in any given flow, allowing to break it down into large- and small-scale structures. With the goal of quantifying different flow regimes and developing computationally-efficient ROMs, POD has been applied to several vascular flow studies, either using numerical CFD data or experimentally-derived velocity fields acquired via Particle Image Velocimetry (PIV). For instance, Kefayati *et al*. [20] used a combination of PIV and POD to study transitional flows in stenosed silicon models; Byrne *et al*. [21] introduced entropy to quantify the flow instability of intracranial aneurysm using POD; Ballarin *et al*. [22] developed a framework for the study of haemodynamics in three-dimensional patient-specific configurations of coronary artery bypass grafts; Chang *et al*. [23] used a POD based ROM to study the flow pattern and the Wall Shear Stress WSS distribution in Abdominal Aortic Aneurysms (AAA). More recently, Buoso *et al*. [24] developed a ROM of blood flow for non-invasive functional evaluation of the pressure drop in coronary artery disease (which computes faster by a factor of about 25 compared to the conventional model), Di Labbio *et al*. [25] compared POD and DMD reconstructions of *in vitro* ventricular flow in a healthy left ventricle and multiple severities of aortic regurgitation, and Han *et al*. [26] applied POD to estimate the flow-induced WSS in computational models of abdominal aortic aneurysm.

An extension of the POD method, Robust Proper Orthogonal Decomposition (RPOD) or Robust Principal Component Analysis (RPCA) has been developed recently to handle noisy or corrupted data, which is often the case for clinical and experimental datasets [12, 27]. However, the application of RPOD algorithms in fluid flows, and especially in the field of physiological flows, is very limited. Some examples are the work by Scherl *et al*. [27] who implemented RPCA filtering to a turbulent channel flow simulation and showed that the coherent flow structures can be extracted from the de-noised low-rank matrix, and the study by Baghaie *et al*. [28] who applied RPCA to filter out the background motion from raw PIV sequences.

In this work, we demonstrate the potential of the RPOD algorithm when compared to traditional POD in an aortic flow. The RPOD method was applied to the patient specific aortic, PIV-derived, flow field described in our previous work [1, 2]. The eigenflows generated are compared to those derived by POD applied to the same data set as well as CFD derived flow fields for the same geometry. ROMs are then successfully used to reconstruct the original flow fields and their potential for personalised hemodynamic modelling is discussed.

## 2. Materials and Methods

A schematic of the approach followed in this work is shown in Figure 1. We have previously characterised and fully validated the flow in a patient specific dissected aorta both experimentally (using PIV) and numerically (using CFD) [2, 1] (black part of the figure). These datasets comprise the Full Order Models. The RPCA algorithm was applied to the PIV velocity field to create a de-noised (reduced-rank) dataset, which we call RPCA velocity field **u**(*x, y, t*)_*RPCA–FOM*_ (blue part of the figure.)

**Figure 1:**
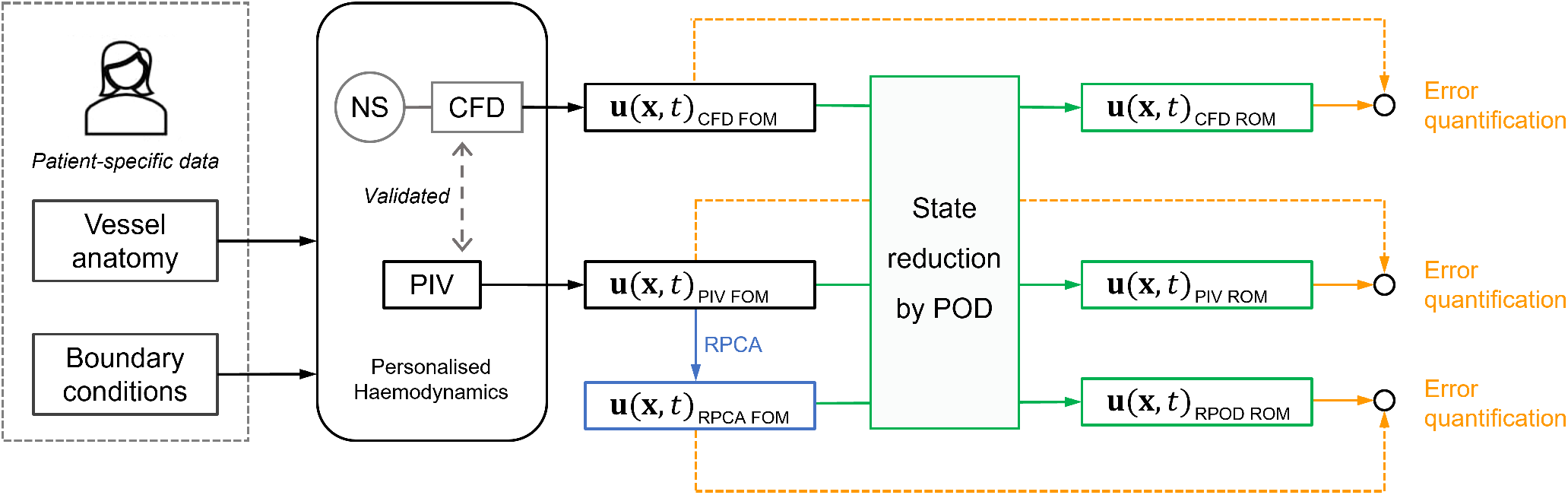
Schematic of the approach followed in this work, comprising four phases. First, the development of Full order models (FOMs), was described in previous works by the authors [1, 2] and led to the experimental PIV model and a computational CFD one (black part of the figure.) In the second phase, RPCA is applied to the PIV velocity field to create a de-noised RPCA velocity field (blue part of the figure.) The third phase involves the creation of ROMs based on the original velocity fields through POD (green part of the figure.) In the last phase, the reconstructed velocity fields were compared to their respective original FOMs, the errors involved were assessed (yellow part of the figure.)

The state reduction of the problem was then achieved by projecting the CFD **u**(*x, y, t*)*_CFD–FOM_*, PIV **u**(*x, y, t*) *_PIV – FOM_* and RPCA velocity fields **u**(*x, y, t*)_RPCA–FOM_ onto POD bases to reduce the dimensionality of the problem (Galerkin projection.) ROMs were identified, ROM-derived flow fields were reconstructed (green part of the figure) and compared to the FOMs. The errors introduced when considering a lower dimensional model were assessed (Yellow part of the figure.)

### 2.1. Patient-specific model

The study is based on clinical data acquired from an adult male with a Type B aortic dissection, a pathology that occurs when a tear in the vessel wall allows blood to flow within the layers of the aorta, leading to the formation of two separate flow-channels, the true and the false lumen. The aortic model was created from the patient CT scans using a semi-automated segmentation tool based on thresholding operations, implemented in ScanIP (Synopsys, Mountain View, CA, USA). It includes one inlet and four outlets: the brachiocephalic trunk (BT), left common carotid (LCC), left subclavian artery (LSA), and descending aorta (DA). A rigid, transparent phantom was manufactured by 3D printing technology (Materialise, Belgium) to enable the flow field measurements described below.

### 2.2. Experimental setup and PIV measurements

The phantom was connected to a custom-made pulsatile flow circuit which comprised a computer controlled pulsatile pump and left ventricle simulator, tunable 3-element Windkessel (WK3) model at each aortic outlet (Figure 2) and an atrial reservoir [29]. The mock loop components were informed by clinical data to reproduce personalised, accurate haemodynamics [2]. A blood mimicking fluid comprising a potassium thiocyanate (KSCN) water solution (63% by weight) was used, matching the refractive index of the phantom. Patient specific flow and pressure waveforms were introduced at the inlets and outlets of the aortic model as illustrated in [1, 2] To perform the PIV measurements, the flow was seeded with fluorescent microparticles with a mean diameter of 10 *μm*, injected into the flow upstream of the phantom and allowed to disperse uniformly within the aortic model. The flow was illuminated by a pulsed Nd:YAG laser (Litron Lasers, Bernoulli, UK) emitting 532 nm wavelength light. Particle image pairs were acquired with a CCD camera (Imperx, USA) at a sampling rate of 22 Hz (the pulsatile flow has a frequency of 1.2 Hz) with a resolution of 4000 × 3000 pixels with a time interval of 1ms, for 10 cardiac cycles. Velocity fields were generated using the Fast Fourier transform based cross-correlation algorithm, implemented with a three-pass technique starting with an interrogation area of 64 × 64 pixels and ending with an area of 32 × 32 pixels, overlapping by 50%. Lastly, post-processing was performed using custom developed MATLAB (MathWorks Inc., USA) functions.

**Figure 2:**
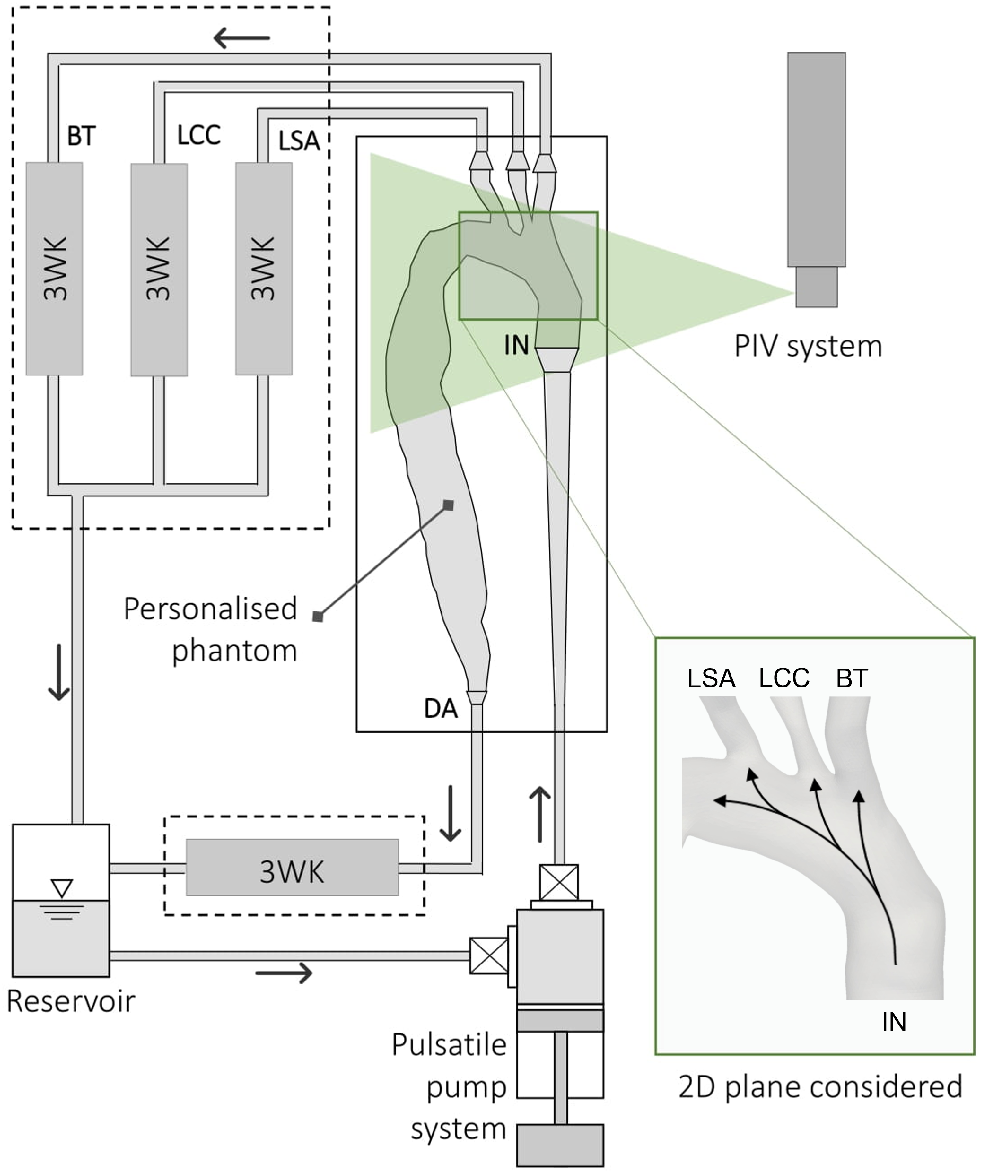
Schematic of the experimental setup. The rig comprises a pulsatile pump system to provide the patient-specific inlet flow rate into the aortic phantom; four 3-elements Windkessel models (3WKs) – one for each of the outlets: brachiocephalic trunk (BT), left common carotid (LCC), left subclavian artery (LSA), and descending aorta (DA) – and an aortic reservoir. The 2D plane where the PIV acquisitions considered in this work were performed is also represented. Superimposed arrows qualitatively indicate the direction of the flow during systole.

Details about the components of the mock circulatory loop and the experimental procedures can be found in our previous work [2, 29]. Here, the PIV-derived velocities obtained on a cross-sectional plane of the aortic arch (shown in Figure 2) were used.

### 2.3. PIV data enhancement by RPCA

Robust Principal Component Analysis (RPCA) or Robust Proper Orthogonal Decomposition (RPOD) is an extension of PCA or POD that can separate an original data matrix (**U**) into a sparse matrix of noise (**S**) and a low-rank matrix that contains *coherent* information (**L**):

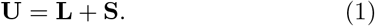

RPCA is known for its ability to handle noisy data [27, 12, 30, 11] - unlike POD that is sensitive to non-Gaussian noise and outliers- and is thus a good candidate to enhance the PIV data in this study prior to further analysis.

RCPA was implemented using the numerical method found in Brunton and Kutz [11]. This entails solving numerically the following minimisation problem, subject to the constraint of Equation 1:

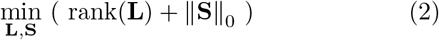

where ||**S**||_0_ represents the zero norm of **S**, which means the summation of non-zero elements in **S**. A convex relaxation 2 form of the problem is used:

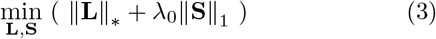

where ||**L**||* represents the nuclear norm of L - which is the summation of all the singular values of **L** - and ||**S**||_1_ denotes the first norm of **S**. λ_0_ is a hyperpa rameter introduced as part of the relaxation given by 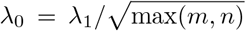 where *m* × *n* is the dimension of **U** and λ_1_ = 1 in the original paper [30]. Scherl *et al*. [27] suggested that λ_1_ can also be used as a tuning parameter: high λ_1_ yields high sparsity of **S**, and low λ_1_ gives low rank of **L**. For simplicity, λ_1_ is kept at 1 in this study. The Augmented Lagrange Multiplier (ALM) algorithm together with the Alternating Direction Method (ADM) [27, 30, 11] were used to solve Equation3.

It should be noted that RPCA is used here as a denoising tool to improve the PIV velocity field and thus is not applied to the CFD data^2^. The term RPCA velocity field is used to denote the PIV velocity field that has been de-noised by the RPCA process, whereas the term RPOD refers to the implementation of POD after RPCA.

### 2.4. Numerical simulation

A detailed description of the numerical procedures for this flow can be found in [1]. ANSYS-CFX 19.0 (ANSYS, USA) was used to solve the 3D incompressible unsteady Reynolds-averaged Navier–Stokes (RANS) and continuity equations, simulating the flow conditions of the PIV experiment. The vessel geometry is the same as the one used to make the phantom, with the walls assumed rigid. All boundary conditions are set to match those from the experiments. The experimental inlet flow rate waveform is imposed with a flat velocity profile. 3WKs model is coupled at the outlets. The fluid is assumed to be Newtonian, with the properties of the KSCN solution. The shear stress transport (SST) turbulence model is chosen with the turbulence intensity of 1% prescribed at the inlet. The simulations run for 3 cardiac cycles and the first two cycles were excluded from the analysis as they contain transient behaviour influenced by the initial conditions and numerical setup. The flow field on the 2D plane aortic arch plane corresponding to the experimental one used here was exported to MATLAB via CFD-Post (ANSYS) for the data reduction.

### 2.5. Proper Orthogonal Decomposition

The POD method decomposes the flow into a set of *modes* arranged depending on their energy content. The higher energy modes represent the coherent structures in the flow; as a result POD has been applied widely to turbulent flows to extract dominant structures. A detailed description of POD can be found in Berkooz *et al*. [31] and in the textbook by Brunton and Kutz [11] and only a brief overview is provided here.

POD is commonly implemented using the method of *snapshots* which is more computationally efficient. Consider a 2D velocity field of *n* = *N_x_* × *N_y_* spatial velocity vectors (*u,v*) on a Cartesian grid *x, y* and a total number of *m* instantaneous velocity fields or *snapshots*. POD decomposes the fluctuating part of the velocity field **u**′(*x,y,t*) into a set of spatial functions **Φ**_*i*_(*x, y*), called the *POD modes*, weighted by time-dependent coefficients *a_i_*(*t*) so that:

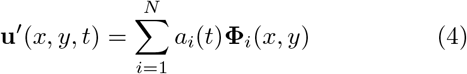

where *i* denotes the mode number.^3^

To perform the decomposition (Equation 4), the time-averaged velocity 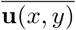 is first subtracted from each instantaneous velocity field, obtaining a set of *m* fluctuating velocity fields **u**′(*x,y,t*). The dataset is then rearranged in a 2*n* × *m snapshot matrix* **U**:

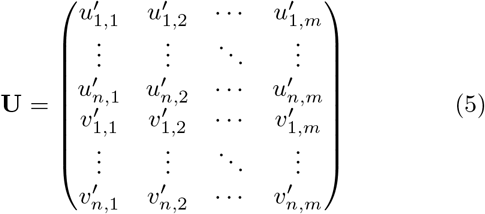

and Singular Value Decomposition (SVD) is applied:

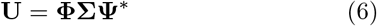

where **Φ** and **Ψ** are the left and right singular vectors of **U**, respectively and the singular matrix (**Σ**) contains the singular values (*σ_i_*) of **U**. The latter rank in descending order, and are directly linked to the portion of kinetic energy (*λ_i_*) contained in the POD modes (**Φ**_*i*_), i.e. 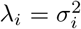. Therefore, POD modes are ranked according to their energy content, with the first mode having the highest energy, and the last the lowest. The total energy captured by the POD modes is defined as the sum of all λ_*i*_:

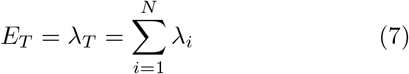

while the energy fraction of the *i*th mode *E_i_*, is defined as

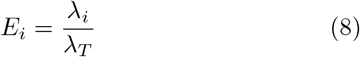

The temporal POD coefficients can be obtained by projecting **U** onto the basis of **Φ**_*i*_:

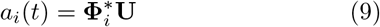

Once the most energetic modes are identified, the velocity field can be reconstructed from the POD modes and the mean velocity 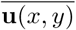 as:

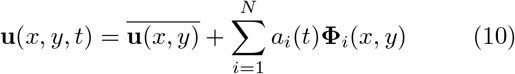

The total number of POD modes (*N*) is the rank of **U**, which is usually equal to the number of snapshots considered. For the PIV data in this study, 10 cardiac cycles with 18 instants per cycle were available, leading to 180 modes in total. For the CFD data, the flow field considered consists of only one cardiac cycle with 165 snapshots, resulting in 165 modes in total. The RPCA velocity field, on the other hand, consists of 180 snapshots since it was generated from PIV data, but only 35 modes. This is because the RPCA process seeks to obtain a low-rank representation of the original data.

### 2.6. State reduction and aortic flow analysis

To generate a low dimensional representation of the aortic flow under consideration, Equation 10 was applied to the most energetic modes *r* identified by the POD. The reconstruction error was estimated from:

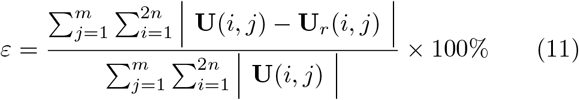

where **U**_*r*_ is the reconstructed snapshot matrix:

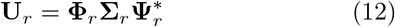

with **Φ**_*r*_, **Σ**_*r*_, and **Ψ**_*r*_ are the truncated versions of **Φ**, **Σ**, and **Ψ**, respectively.

The POD spatial structures and temporal coefficients were used to characterise specific flow features - *coherent structures* - in the pulsatile, aortic dissection flow, separating the periodic and random fluctuating structures from the mean flow. The spatial structures **Φ**_*i*_ for the relevant modes were analysed plotting the velocity fields. Then, the temporal characteristics of the flow were investigated by analysing the temporal coefficients of the most energetic POD modes in both the time and frequency domains. The relation between the first coefficients *a_i_* was investigated by plotting the space 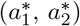, calculated as:^4^

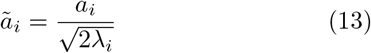

and normalise as

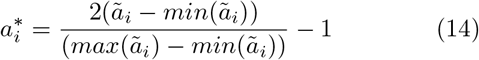

Lastly, flow field reconstructions from the ROMs were performed according to Equation 10. First **u**(*x,y,t*) was reconstructed using all the modes to verify the accuracy of the mathematical calculations. Then, they were reconstructed using only a selected number of modes (i.e. **Φ**1-2, **Φ**1-5, and **Φ**1-10) and the solution was compared to the original velocity field, at different instants of the cardiac cycle, to quantify the differences.

## 3. Results and Discussion

### 3.1. Kinetic energy distribution

Table 1 lists the cumulative kinetic energy contents of the POD and RPOD modes derived from the PIV data (PIV POD, PIV RPOD respectively) compared to those derived from the CFD data (CFD POD). While more than 90% of the kinetic energy is reached within the first 2 modes for the PIV RPOD and CFD POD data, it takes 10 modes for the PIV POD to capture that amount of energy. This is not surprising as the PIV data are subject to measurement noise. The latter is more pronounced during diastole where the measured velocities are low and comparable to experimental errors, found in the order of 5% [2]. The RPCA process denoises the PIV data resulting in higher cumulative energy in the first two and ten modes compared to that from PIV data alone. The energy content of the POD modes becomes closer to that derived from the CFD data which are highly resolved and free from cycle-to-cycle variations and measurement noise.

**Table 1:** Percentage of Kinetic energy captured and reconstruction error of different groups of POD modes calculated from PIV velocity field, RPCA velocity field, and CFD velocity field

| *Modes* | PIV POD |  | PIV RPOD |  | CFD POD |  |
| --- | --- | --- | --- | --- | --- | --- |
|  | Energy(%) | Error(%) | Energy(%) | Error(%) | Energy(%) | Error(%) |
| 1-2 | 86.33 | 34.18 | 98.15 | 13.10 | 95.52 | 19.54 |
| 1-5 | 89.57 | 30.16 | 99.37 | 7.85 | 98.77 | 10.18 |
| 1-10 | 91.72 | 27.06 | 99.78 | 4.60 | 99.72 | 4.81 |
| all | $\sim 100$ | $\sim 0$ | $\sim 100$ | $\sim 0$ | $\sim 100$ | $\sim 0$ |

### 3.2. POD structures and temporal coefficients

Selected POD spatial structures (**Φ**1-3) are shown in Figure **??** compared with the mean flow. The eigenflows correspoding to the first POD mode (**Φ**1) show organised motion in the same direction as the mean flow which reflects the flow at the peak systolic phase [2]. The eigenflows extracted from PIV POD and PIV RPOD appear to be almost identical as they represent the most energetic flow features; they slightly differ from the CFD POD ones due to the differences between the measured and computed velocity fields discussed in our previous work [1].

Modes 2 and 3 (**Φ**2-3) exhibit more complex flow patterns characterised by re-circulation regions. As for **Φ**2, a high magnitude region can be seen at the inner side of the arch which reflects the flow pattern in diastole.

The temporal coefficients *(a_i_)* of the first three POD modes are shown in Figure 4 in the time (left column) and frequency domain (right column), respectively. The coefficients exhibit periodic characteristics in agreement with the literature *et al*. [20]. The first temporal coefficient essentially reflects the shape of the patient-specific inlet flow waveform manifesting with a dominant peak in the spectra at the frequency of the cardiac cycle (f = 73.2 bpm = 1.22 Hz). As the number of modes increases, the temporal coefficients show more complicated patterns characterised by higher frequency oscillations. A second harmonic (2.44 Hz) is evident on the frequency spectra of the experimentally derived modes. This behaviour may be related to velocity fluctuations due to transitional flow. This behaviour is absent from the numerical POD coefficients which consist of one single cardiac cycle only, hence oscillating at the cardiac cycle frequency only.

**Figure 3:**
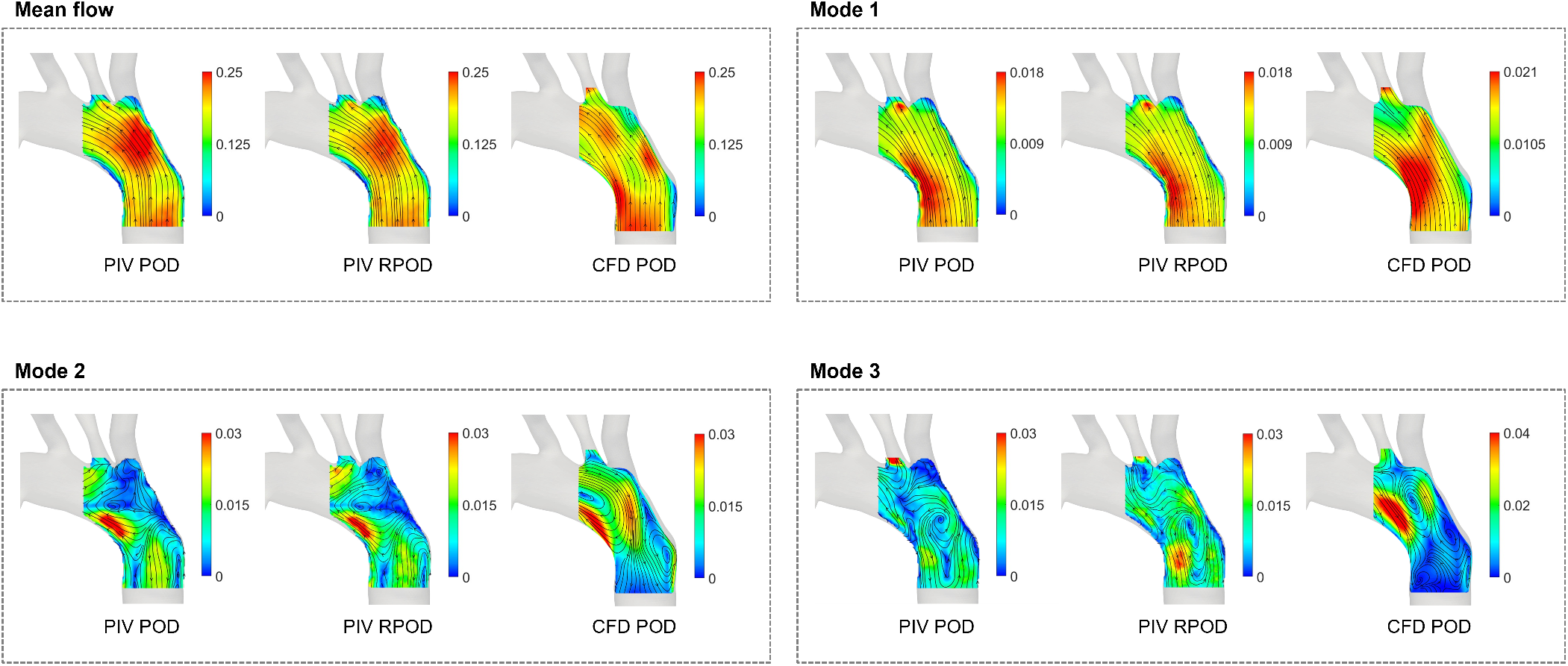
Velocity fields contours of the mean velocity field and those of the first three modes (**Φ**1-3) obtained from the PIV, RPCA, and CFD velocity fields, respectively. Super-imposed streamlines were used for illustration purposes, they do not convey any information on temporal variations.

**Figure 4:**
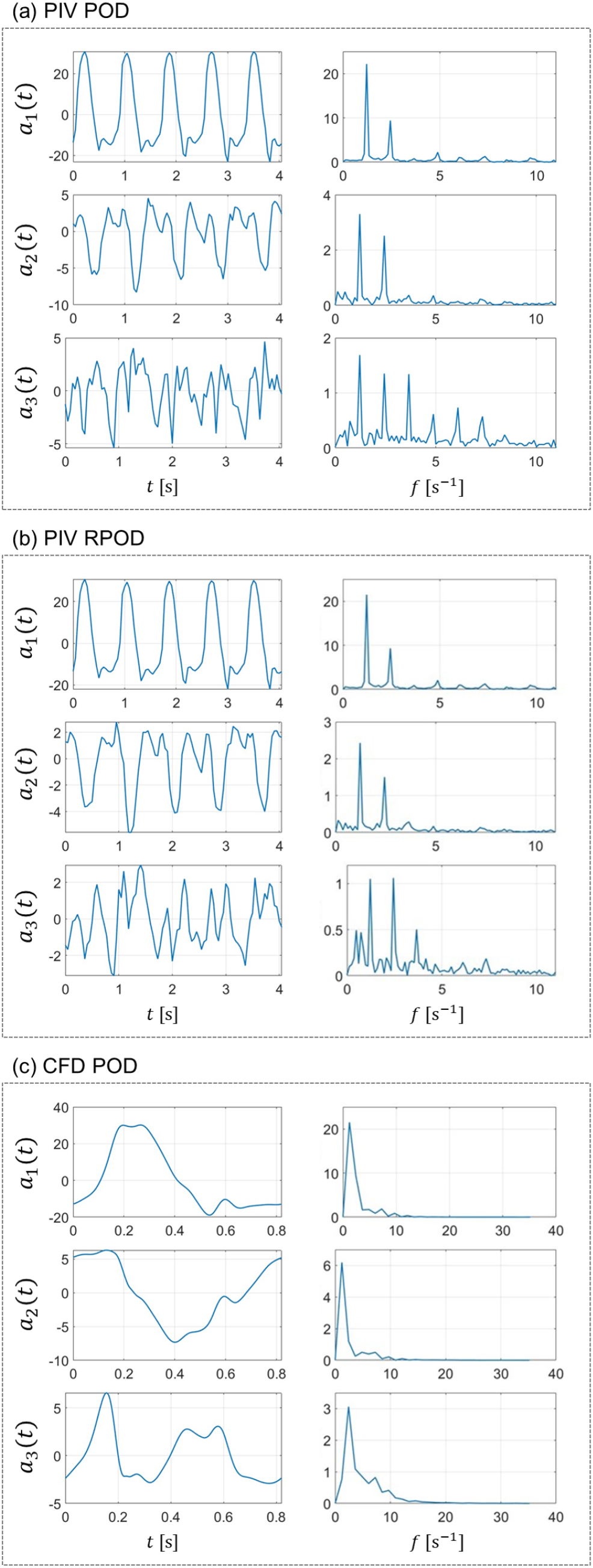
The first three POD temporal coefficients (*a*_1_, *a*_2_, and *a*_3_) from (a) PIV POD, (b) PIV RPOD, and (c) CFD POD presented in (left) time domain and (right) frequency domain.

The normalised temporal coefficients 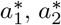 and 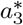, are plotted against each other in Figure 5. The phase-averaged coefficients for the experimental data are also indicated in blue. They exhibit organised, closed-loop structures, indicating periodicity similar to the CFD data.

**Figure 5:**
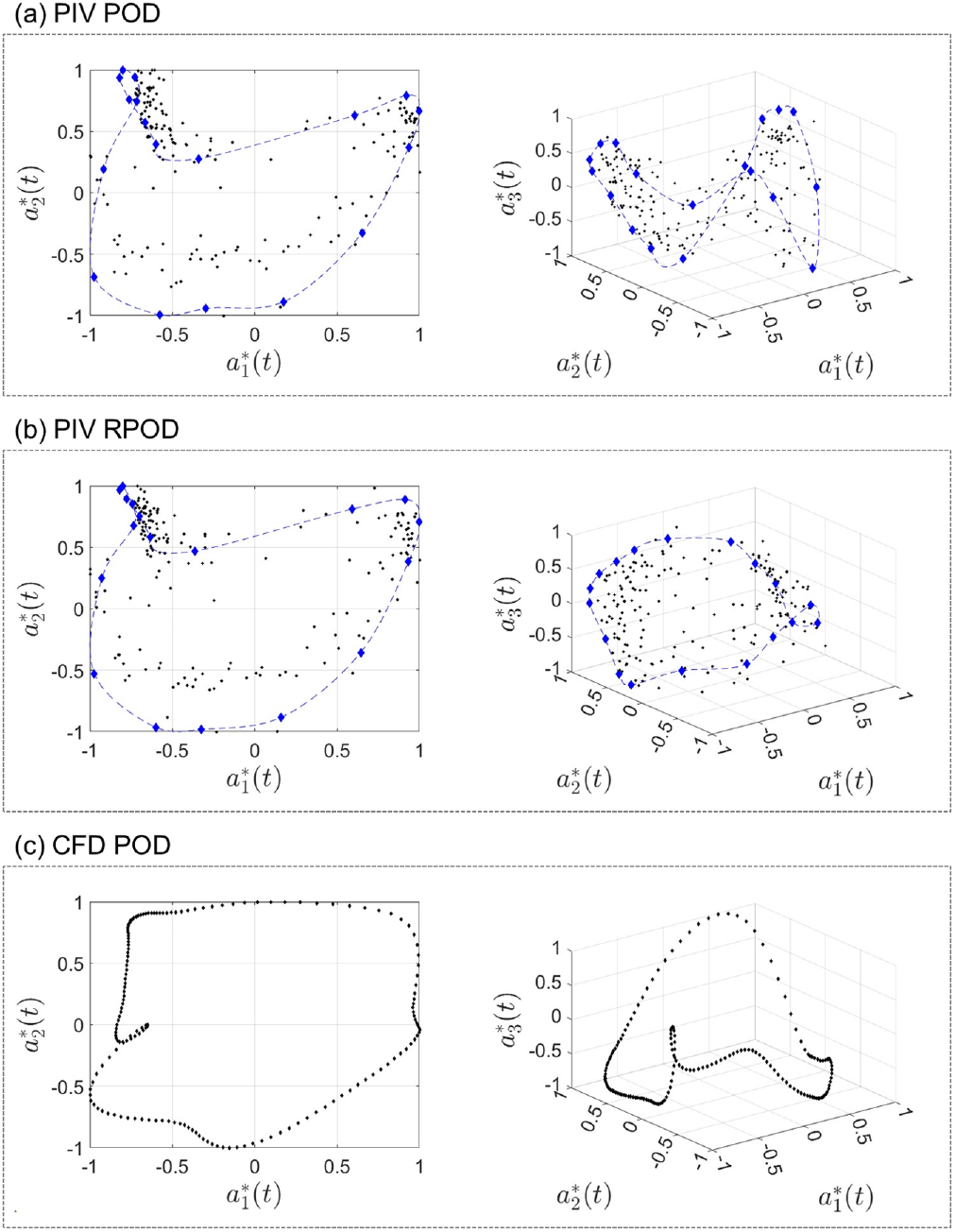
Scatter plots of normalised POD coefficients (left) 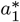 and 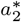 and (right) 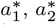 and 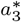 computed from (a) PIV POD, (b) PIV RPOD, and (c) CFD POD. The black points represent the coefficients for all the modes, whilst the blue ones represent the phase-averaged POD modes. The blue lines connect the phase-averaged coefficients for better visualisation.

The 2D plots of 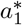 and 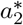 in Figure 5 do not show a clear circular or elliptic pattern, implying that the first two POD modes do not form a pair. The plots also indicate the same behaviour for the first two PIV POD and PIV RPOD coefficients which slightly differs from the CFD POD ones.

An interesting observation arises when investigating the relation amongst the first three coefficients (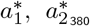 and 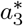) The plots on the right side of Figure 5a-c show a more complex organisation amongst these modes. This behaviour highlights the interdependent relationships and energy transfer between the first three modes and may correspond to energy transfer amongst different periodic structures (with different energy contents and frequency profiles) within the flow field. A similar *‘triadic interaction’* has been reported by Gabelle *et al*. [34] in a stirred tank flow (who attributed the behaviour to non-linear interactions between the modes), and by Lacassagne *at al*. [35] in an oscillating grid flow.

### 3.3. Flow reconstructions

Figures 6–8 show the comparison between the FOMs at three instants of the cardiac cycle (peak systole, deceleration and diastole), and the reconstructed velocity fields of ROMs using **Φ**1-2, **Φ**1-5 and **Φ**1-10. The figure also shows contours of the differences in velocity magnitude between the original and reconstructed flow fields. The reconstruction errors are shown in Table 1. As expected, in all cases, the more POD/RPOD modes are included in the reconstruction, the lower the error.

**Figure 6:**
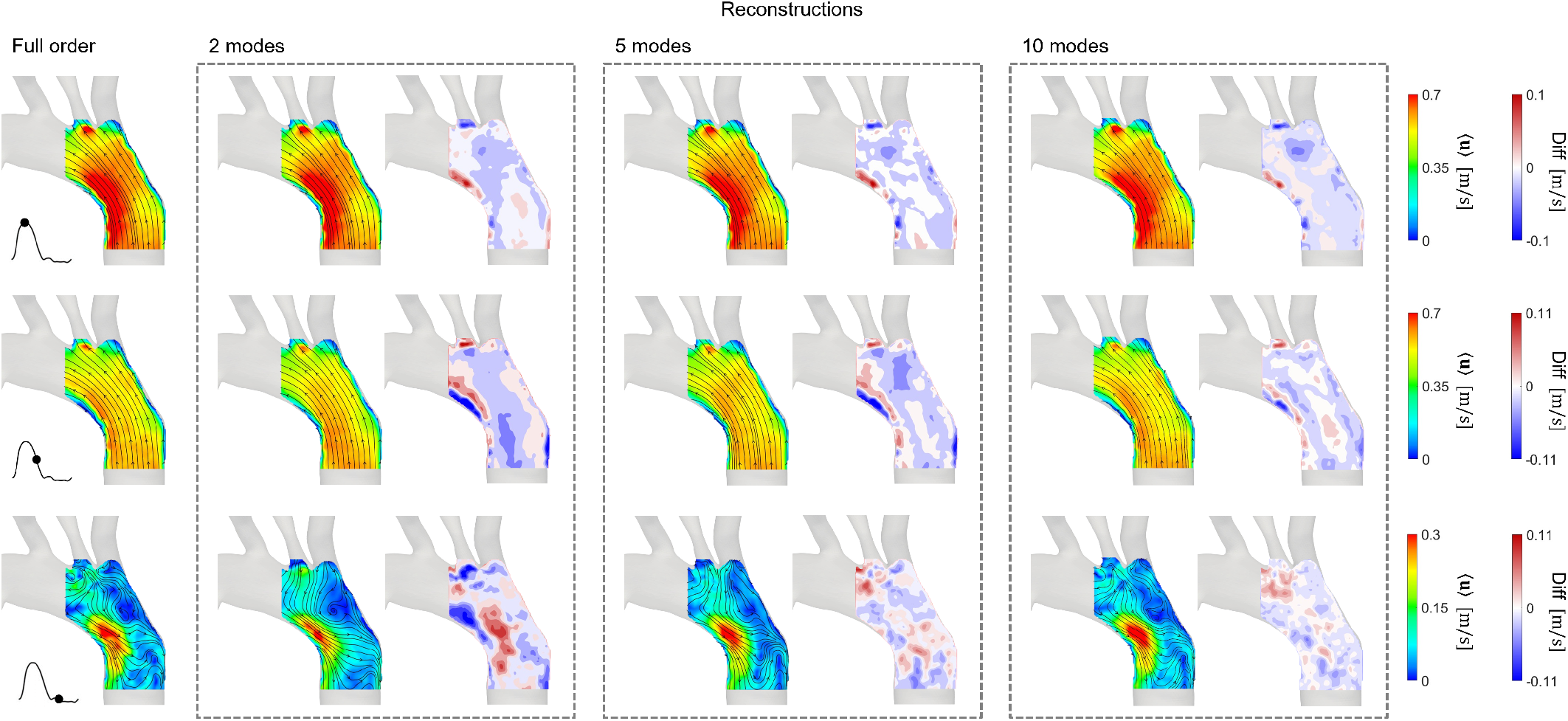
PIV POD, a POD reconstruction of PIV velocity field using **Φ**1-2, **Φ**1-5, and **Φ**1-10. The FOM and reconstructed flow fields are visualised at three instances of the cardiac: peak systole, deceleration, and diastole. Contours showing the difference in velocity magnitude between the FOM and reconstructed flow fields are shown side by side with the reconstructed velocity field.

**Figure 7:**
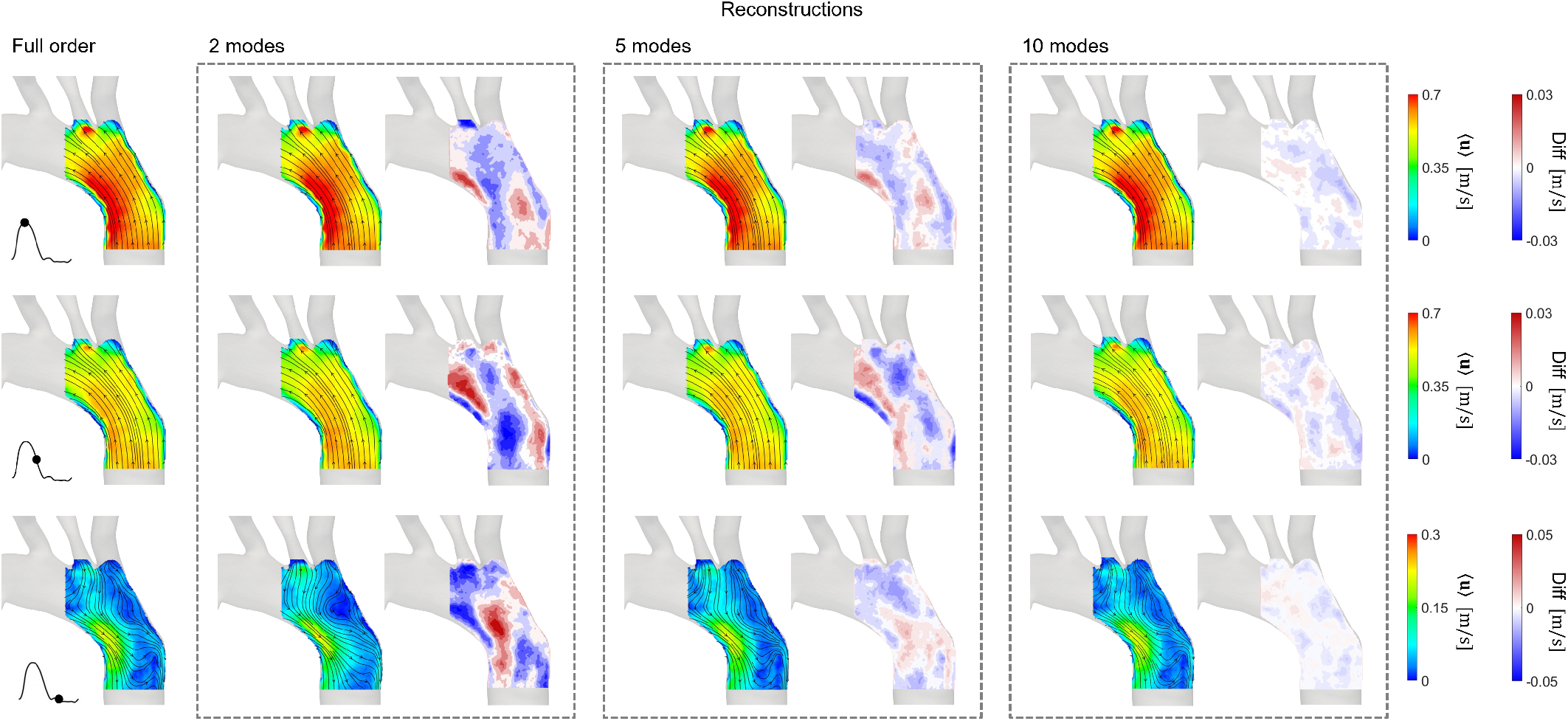
PIV RPOD, a POD reconstruction of RPCA velocity field using **Φ**1-2, **Φ**1-5, and **Φ**1-10. The FOM and reconstructed flow fields are visualised at three instances of the cardiac: peak systole, deceleration, and diastole. Contours showing the difference in velocity magnitude between the FOM and reconstructed flow fields are shown side by side with the reconstructed velocity field.

**Figure 8:**
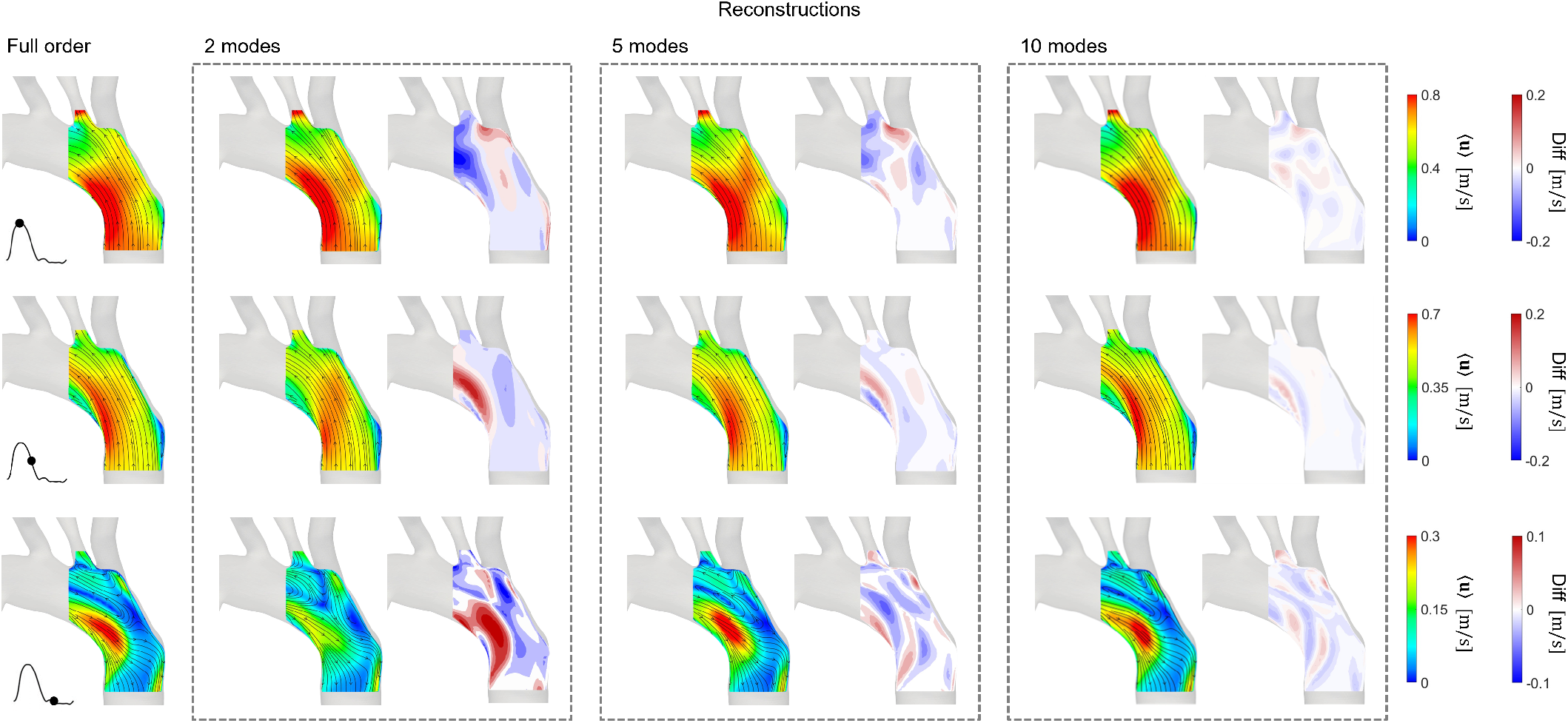
CFD POD, a POD reconstruction of CFD velocity field using **Φ**1-2, **Φ**1-5, and **Φ**1-10. The FOM and reconstructed flow fields are visualised at three instances of the cardiac: peak systole, deceleration, and diastole. Contours showing the difference in velocity magnitude between the FOM and reconstructed flow fields are shown side by side with the reconstructed velocity field.

The reconstructed flow fields from the PIV POD in Figure 6 show that good agreement can be achieved even using the first 2 POD modes at peak systole and in the descending part of the flow curve, where the flow follows relatively organised uni-directional patterns, with some differences occurring in near-wall re-circulation regions. However, more significant discrepancies are observed in diastole, both in terms of velocity magnitude, distribution and the flow directions indicated by the streamlines. Such differences are to be expected because, at diastole, the flow contains smaller vortical structures and a more complex flow field. To fully reconstruct these structures, a higher number of modes should be included. 10 modes appear to be sufficient to reconstruct the flow at diastole accurately. When reconstructing the flow with modes **Φ**1-10, the maximum absolute difference is about 0.09-0.10 m/s and occurs at systole, where the velocity ranges between 0-0.7 m/s.

Similar reconstructed velocity fields are generally obtained from the PIV RPOD in Figure 7. The only noticeable difference is at diastole, where the RPCA velocity fields appear more uniform and exhibit lower velocity magnitudes near the aortic arch compared to PIV derived ones, due to its filtering action. Thus, when using the same number of modes to reconstruct the flow field, PIV RPOD has a much lower reconstruction error than PIV POD. The maximum absolute error when using 10 modes for the flow reconstruction is only 0.007-0.008 m/s, i.e. 10 times lower than that in PIV POD.

Finally, Figure 8 shows that the flow reconstructions from the CFD POD analysis share similar important qualities with the experimental ones; namely that the first 2 modes are able to reconstruct the velocity field accurately at peak systole and deceleration phase, with the errors decreasing rapidly when more modes are included. The difference contour plots show less scatter compared to the PIV derived ones due to high spatial resolution of the numerical data. A maximum difference of around 0.05-0.06 m/s is found when reconstructing the flow fields using 10 modes; this is smaller than the errors in PIV POD but slightly higher than the case of PIV RPOD.

### 3.4. Towards personalised ROMs

The use of the RPCA algorithm in this study successfully filtered out high-frequency noise in the PIV data (see Figure 4), improving the performance of ROMs extracted from the data. However, the disappearance of the high-velocity zone near the aortic arch during diastole in the RPCA flow field strongly suggests that the algorithm may also filter out some important flow features ^5^. This could potentially be due to over-filtering and could be addressed by increasing the value of the tuning parameter λ_1_. However, too high a value of λ_1_ can also lead to the presence of noise in the filtered data. Therefore, the main challenge involved in the application of RPCA is to find the optimal value of λ_1_ that appropriately filters out the unwanted motion while preserving the relevant ones.

The enhancement of *in vitro*, experimental data using RPCA demonstrated here suggests that such methods could potentially be applied to *in vivo* data, such as 4D flow MRI, to improve their quality, making them more amenable to computational modelling and flow reconstruction [36, 37, 38]. In addition, the ability of the RPCA algorithm to effectively clean data may also result in ROMs constructed from RPOD having to include fewer modes, leading to faster computations in their subsequent applications.

This work also demonstrates that it is possible to represent the behaviour of complex, pathological aortic flows using ROMs consisting of only the first few POD/RPOD modes, which shows promise in the development of more computationally efficient models to support clinical decision-making. Examples include the works of Chang *et al*. [23], who developed a computationally efficient ROM to study the flow patterns and the WSS distribution in simplified models of an abdominal aortic aneurysm, and Buoso *et al*. [24], who developed ROMs of blood flow for non-invasive functional evaluation of the pressure drop in coronary artery disease using parameterised POD. ROMs may possess properties that can serve as supplementary haemodynamic indices. For example, by monitoring the temporal evolution of energy distribution, it may be possible to track the progression of some cardiovascular diseases. Moreover, the energy fraction associated with higher-order POD/RPOD modes may contain information that can be used to fine-tune turbulence parameters when modelling vascular flows.

Finally, ROMs can also be combined with the rapidly evolving machine learning tools to allow for *optimization* and *design* in fluid mechanics, moving towards real-time modelling. This would allow, for instance, to study a wide range of parameters for a given vascular pathology (e.g. increasing or decreasing the level of stenosis on coronary disease or coarctations) and analyse the consequences on the flow and pressure fields, which could serve as an initial step to investigate patient-specific pre-interventional options [39, 40].

## 4. Conclusions

The time-dependent flow in an aortic model, measured by PIV, was enhanced by RPCA and decomposed by means of POD to create ROMs. The decomposed flows were compared against those from numerical data obtained for the same patient-specific conditions. The first two modes derived from RPOD capture more than 90 per cent of the kinetic energy similar to the experimental ones, bringing the PIV data closer to the CFD ones in terms of energy content and reconstruction errors.

The large and small-scale structures within the flow, corresponding to more or less energetic modes, were evaluated and described by means of POD/RPOD spatial structures and POD/RPOD temporal coefficients. By combining only the most energetic modes to represent the flow, it was shown that complex, time-dependent haemodynamic data can be represented with simpler low-dimensional models based on a small number of spatial modes. This combined with the strong reconstruction performance of RPOD, illustrates the potential of the approach to enhance the quality of measurements and to develop more computationally efficient models for clinical application.

## Acknowledgements

This project was supported by the Wellcome/EPSRC Centre for Interventional and Surgical Sciences (WEISS) (203145Z/16/Z); the EPSRC Transformative Healthcare Technologies grant PIONEER, EP/W00481X/1; the Department of Mechanical Engineering, University College London; and the British Heart Foundation grant VIRTUOSO (FS/15/22/31356). The authors would also like to thank Dr. Tom Lacassagne and Dr. Andrea Ducci for their constructive feedback.

## Footnotes

1 POD is mathematically equivalent to the popular Principal Component Analysis (PCA) [10]

2 The cleanness of the CFD velocity field relative to that of the PIV velocity field is discussed in Section 3.1

3 Note that **u**′ in Equation 4 includes both the periodic variation around the mean and the turbulence fluctuation.

4 This approach has been used in different studies, for example see [32] and [33].

5 The high-velocity zone appears in both the PIV and the CFD velocity fields, thus it is likely part of the dominant flow features

